# Weakly-Coupled Oscillators with Long-Distance Correlation as a Model of Human Atrial Fibrillation

**DOI:** 10.1101/2021.01.09.425808

**Authors:** Shahriar Iravanian

## Abstract

Atrial Fibrillation (AF) is a chronic, progressive, and heterogeneous disease, which exhibits irregular and chaotic atrial electrical activity. The exact mechanisms of AF have remained elusive. The complexity of the atrial signals, irregular period, and the non-local nature of AF make it very difficult to characterize its spatiotemporal organization. This paper presents a set of data-processing tools to find and evaluate the intracardiac signals in AF based on synchronization theory. Specifically, a graph-theoretical algorithm is developed to prune spurious detections of atrial complexes (spikes). The processing pipeline was applied to 10 intracardiac recordings obtained during ablation procedures in patients with paroxysmal or persistent AF. The AF cases were classified into three main types. Type I (n=3) is defined as the presence of a single driver that dominates both atria and is consistent with the mother rotor hypothesis. The majority of the cases had more than one driver and were classified as types II (n=4) and III (n=3). The drivers, i.e., the interacting organized areas, in type II have significantly different peak frequencies and are only weakly coupled. The drivers in type III have close frequencies, are moderately interacting, and show evidence of transient intermittent phase-locking (intermittency). In addition, a long-distance correlation between channels from the left and right atria was observed. The theory of synchronization is a useful conceptual framework to process and analyze AF and can provide mechanistic insight into AF and the effect of various interventions.

## II. INTRODUCTION

The exact mechanisms responsible for the initiation and perpetuation of atrial fibrillation (AF) have remained elusive.^1^ AF is the most common sustained cardiac arrhythmia, and has been the subject of thousands of experimental, theoretical, modeling, and clinical studies since its recognition in the early twentieth century. Despite all these efforts, there are still major gaps in our knowledge and understanding of how AF is triggered and sustained.

The pioneering modeling works of Moe in the early 1960s proposed multi-wavelet reentry as a mechanism for AF.^2^ Allessie et al, in a series of seminal papers, confirmed the presence of multiple-wavelets during AF in rabbit hearts.^3,4^ Winfree recognized spiral waves as a major organizing feature in two-dimensional excitable media.^5^ Finally, Jalife et al experimentally detected spiral waves in AF.^6^

A major unanswered question is whether AF is sustained by a single reentrant spiral wave (called a *mother rotor*) or whether multiple interacting rotors are responsible for AF.^3,6,7^ In this paper, *driver* is used as a generic term to refer to autonomous and temporally-persistent units of atrial activity during AF. The underlying mechanism of a driver is usually a reentrant, anchored, or meandering, spiral wave. The term rotor is used interchangeably with a spiral wave.

Drivers are the building blocks of AF; however, the spatial organization of these drivers is not well understood. A related question is the effective correlation range during AF. While short-range correlation (1-2 cm) is well-established,^8^ whether points farther than a few centimeters affect each other is unknown. In previous experimental and modeling studies, we postulated that the transition from chaotic and irregular AF to regular flutter (stable, single reentry) in the course of ablation is a type of phase transition and is heralded by a sudden increase in the correlation range.^9,10^

The details of AF mechanisms and dynamics affect the ways ablation are done to control AF. Nowadays, ablation procedures are the mainstay of AF management.^11^ During ablation, an electrophysiologist induces tissue damage at strategic locations in the atrial wall (especially in the left atrium) to stop the AF-initiating triggers originating from pulmonary veins and to disrupt the circuits of the reentrant waves. A successful targeted ablation requires the knowledge of the number and location of the putative spiral waves.^12,13^ As the result of our knowledge gaps and sub-optimal clinical efficacy,^14^ targeted ablation is rarely performed anymore, and most operators settle for a static set of per-determined ablation lesions.

One major obstacle to study AF is the lack of a suitable animal model.^15^ AF is a chronic, slowly-progressive disease and it is uncertain that an animal model that causes AF within days to weeks can capture all the essential characteristics of human AF. Therefore, the best source of information about AF has remained human AF patients. Previously, we reported on the spatiotemporal dynamics of AF based on signals recorded from a 20-pole catheter during ablation of persistent AF.^9,16^ In the current study, the data was the unperturbed (without ongoing ablation) signals recorded using multiple catheters covering both atria. This dataset is complementary to the one we had used previously, as a trade off between temporal coverage for better spatial coverage. An ideal dataset would have both extended temporal and dense and wide spatial coverage; however, safety and ethical considerations preclude collecting such data from patients. Therefore, the focus of the current paper is on the analytic tools developed to analyze the available dataset to gain mechanistic insight about AF. Especially, the graph-theoretical algorithm described in the Spike Pruning section significantly improved the accuracy the data processing pipeline and made it possible to extract meaningful correlations from noisy signals.

## III. METHODS

### A. Data Collection

The data were collected retrospectively from AF ablation procedures. The data collection protocol was approved by the Emory University Institutional Review Board. The data is comprised of intracardiac electrograms recorded with the help of three catheters placed in the right and left atria of patients undergoing AF ablation. The first catheter was a 20-pole (10 bipolar pairs) catheter that covered the tricuspid annular region of the right and the proximal coronary sinus area of the left atrium. The second catheter was one of the following mapping catheters placed in the left atrium and parked at or near one of the pulmonary veins: a circular mapping catheter (20 poles, 10 bipolar pairs), a flat two-dimensional grid (Abbott HD Grid, 4×4 electrodes, set up as 24 bipolar pairs between adjacent electrodes), or a multi-spline mapping catheter (Biosense Webster Penta-Ray, set up as 10 bipolar pairs). An ablation catheter was also placed in a pulmonary vein contra-lateral to the mapping catheter. Each data segment was at least five minutes of unperturbed AF and was recorded before the start of ablation with no catheter movements throughout the recording. The signals were band-passed filtered at 30-500 Hz, digitalized at 12 bits of resolution, sampled at 977 Hz, and were downloaded for offline processing.

### B. Spike Detection

Figure 1A shows a typical intracardiac recording during AF. As a result of high-pass filtering, the signal is made of distinct spikes, corresponding to the local upstroke of the action potentials and lacks repolarization information. The first step in processing the data is the detection of the spikes. Because of significant beat-to-beat variability during AF, caused by the chaotic nature of AF, variable wavefront direction, and heart motion, spikes in the same channel can have different shapes and amplitudes. Therefore, the spike detection algorithm needs to be robust against these artifacts. A peak-detection algorithm based on continuous wavelet transform, which was originally designed to analyze mass spectrograms, was adapted to detect atrial complexes during AF.^17^ The main tuning parameter for the algorithm is the widths of the wavelets used to smooth the signal. The algorithm works very well for highly periodic signals during atrial flutter but tends to miss some spikes and over-detect spurious ones during AF. The solution is to run the algorithm with a wide range of wavelet widths (10-30 ms) to ensure a low under-detection rate while accepting over-detection (Figure 1B). The spurious peaks are then filtered out in the next step (Figure 1C, see Spike Pruning).

**FIG. 1.**
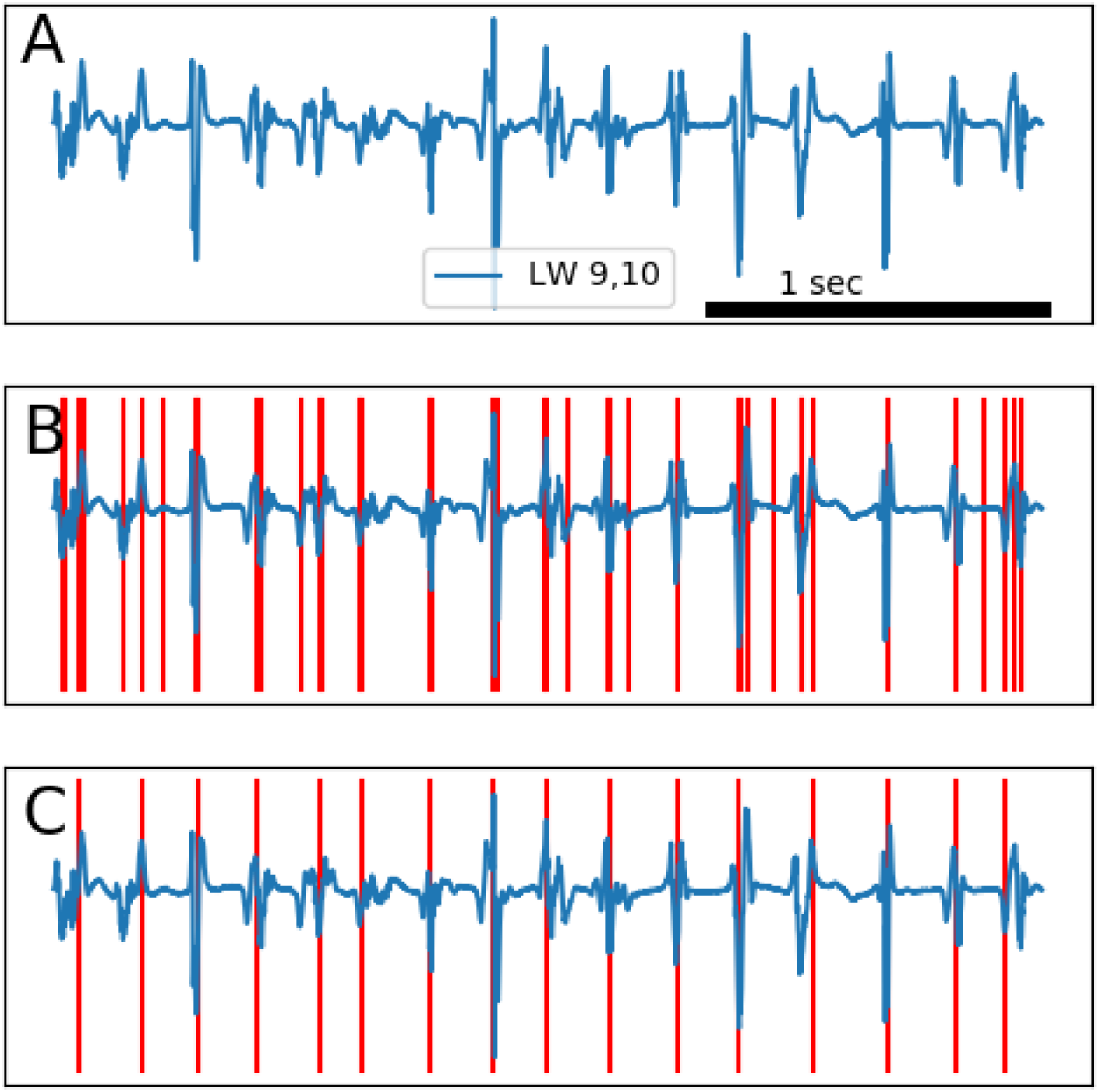
Representative intracardiac electrograms during AF. (a) Raw signals (band-pass filtered at 30-500 Hz). (b) The same signal as (a) with the super-imposed candidate spikes (red lines), detected using the continuous wavelet transform based algorithm. (c) True spikes after the pruning pass.

### C. Spike Pruning

Let *t*(*i*), for *i* = 1 *… N*, be the timing of the *i*th spike detected in the previous stage. The set of *t*(*i*) contains both true and false spikes. Our goal is to filter out the false detections and find the most probable (in the maximum likelihood sense) subset of the spikes, given the expected statistics of the intracardiac signals. Let *D* = (*d*_1_,…, *d*_*j*_,…, *d*_*M*_), for 1 *≤ d*_*j*_ ≤ *N* and *d*_*j*_ < *d*_*j*+1_, be an ordered list of *M* indices. We want to find *D* such that *t*(*d*_*j*_) marks a true spike. We assume that,

1. Cycle length, *T*_*i*_ = *t*(*i*)*−t*(*i−*1), is a random variable drawn from a normal distribution 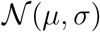, where *μ* and *σ* are the mean and standard deviation, respectively. The median of *T*_*i*_s is a robust estimation of *μ*. As we will see, *σ* will cancel out and its value is not needed.
2. The consecutive cycle lengths, *T*_*i*_ and *T*_*i*+1_, are drawn independently from 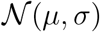, i.e., *T*_*i*_ is an independent and identically distributed random variable. This assumption is not strictly true, but given the short mixing time of AF (less than a second), is a reasonable approximation.

Based on our assumptions, we can assign a probability to a given *D* as

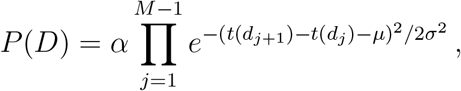

where *α* is a normalizing factor. In fact, for simplicity, 2*σ*^2^ in the exponent can be absorbed into *α*. Therefore,

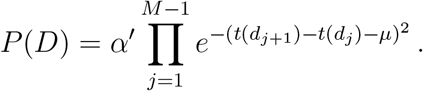

We want to find the subset that maximizes this probability, or equivalently, minimizes the negative of its logarithm,

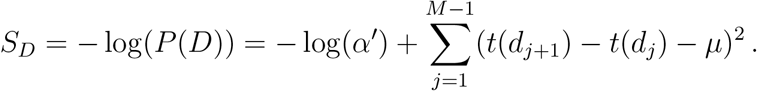

The normalizing factor, *α*^’^, depends only on *σ* and can be ignored. We need to find *D* that minimizes the summation term.

Let us look at a simple example. Assume we try to find the best solution for a short segment *t*(1)*, t*(2)*, t*(3)*, t*(4), where *t*(1) and *t*(4) are already accepted as true spikes. There are four possibilities. First, *t*(2) and *t*(3) are both true spikes, then we have *D* = (1, 2, 3, 4). Therefore, *S*_*D*_ = *S*_(1,2,3,4)_ = (*t*(2) *− t*(1) *− μ*)^2^ + (*t*(3) *− t*(2) *− μ*)^2^ + (*t*(4) *− t*(3) *− μ*)^2^. The next option is *t*(2) a true spike and *t*(3) a false one; hence, *D* = (1, 2, 4) and *S*_(1,2,4)_ = (*t*(2) *−t*(1) *−μ*)^2^ +(*t*(4) *−t*(2) *−μ*)^2^. Similarly, *S*_(1,3,4)_ = (*t*(3) *−t*(1) *−μ*)^2^ +(*t*(4) *−t*(3) *−μ*)^2^ and *S*_(1,4)_ = (*t*(4) *− t*(1) *− μ*)^2^. Depending on which *S* is smaller, we can choose the right option. Now, let’s assume we have a segment *t*(*i*),*t*(*i* + 1),*t*(*i* + 2),*t*(*i* + 3) embedded inside a longer sequence. In this situation, we cannot simply assume that *t*(*i*) and *t*(*i* + 3) are true spikes. Instead, whether *t*(*i* + 1) or *t*(*i* + 2) are accepted affects the acceptance of *t*(*i*) and *t*(*i*+3) and vice versa. In general, *D* is picked from 2^*N*^ possibilites and a brute-force approach is not feasible. The solution is to cast the optimization problem as a graph problem, such that the optimized solution corresponds to the shorted path on the graph. The resulting graph problem can be solved using a standard shortest path algorithm (e.g., the Dijkstra’s algorithm).

Figure 2 depicts the problem graph, *G* = (*V, E*), where *G* is a directed weighted graph and *V* is the set of *N* vertices, one for each *t*(*i*). For each pair of vertices *i* and *j*, such that *i < j*, we add a directed edge *e* with weight (*t*(*j*) *− t*(*i*) *− μ*)^2^ to *E*. For simplicity, we assume that *t*(1) and *t*(*N*) are always accepted (it is possible to relax this restriction by adding dummy nodes). The shortest path from vertex 1 to *N* defines the *D* that minimizes *S*_*D*_; therefore, we can accept the nodes traversed by the shortest path as true spikes (Figure 2B).

**FIG. 2.**
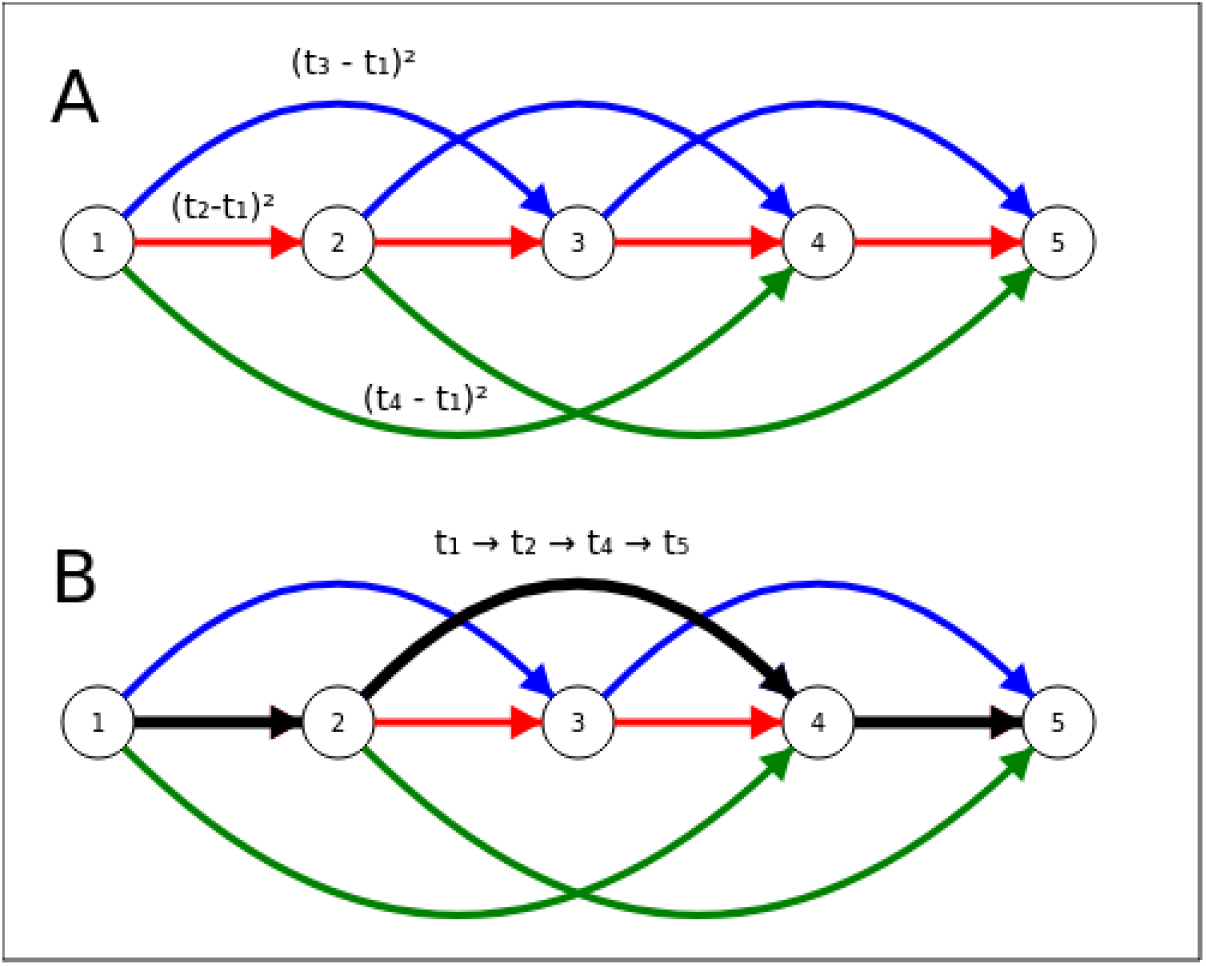
The schematic diagram of the graph-based pruning algorithm. (a) The spikes graph for 5 nodes *t*(1)*,…, t*(5). Some edges are marked with the corresponding weight. Note that the edge from 1 to 5 is skipped for clarity. (b) An example shortest path (the black line). For this path, the true spikes are *t*(1) *→ t*(2) *→ t*(4) *→ t*(5).

### D. Phase Transformation

After we find the true spikes for each channel, the next step is to convert the spike data into a phase signal. This is done as an analogous technique to the Poincare maps by linear interpolation between two consecutive spikes and assigning a phase value between 0 to 2*π* to the intervening points,^**?**^

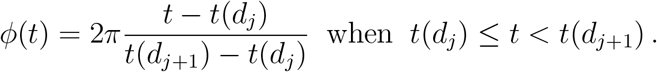

This method is sensitive to false spike detection, which is why the spike pruning algorithm is needed. Let *φ*_1_(*t*) and *φ*_2_(*t*) be the phase signals calculated from two different channels. In this paper, the main results are based on the comparison of *φ*_1_ and *φ*_2_ using recurrent plots as a visual guide to detect subtle correlation between channels and stroboscopic histograms to calculate the statistical significance of the presumptive association.

Figure 3A shows a typical recurrent plot. The range 0 to 2*π* is divided into 20 bins, *φ*_1_ and *φ*_2_ are sampled every 10 ms, and the respective bin numbers are added to form a 20×20 2D histogram, which is visualized as a recurrence map. Lines parallel to the principal diagonal are indicative of 1:1 correlation between *φ*_1_ and *φ*_2_. Higher order relationships (e.g., 2:1 and 3:2) manifest as lines with different slopes than 1.

**FIG. 3.**
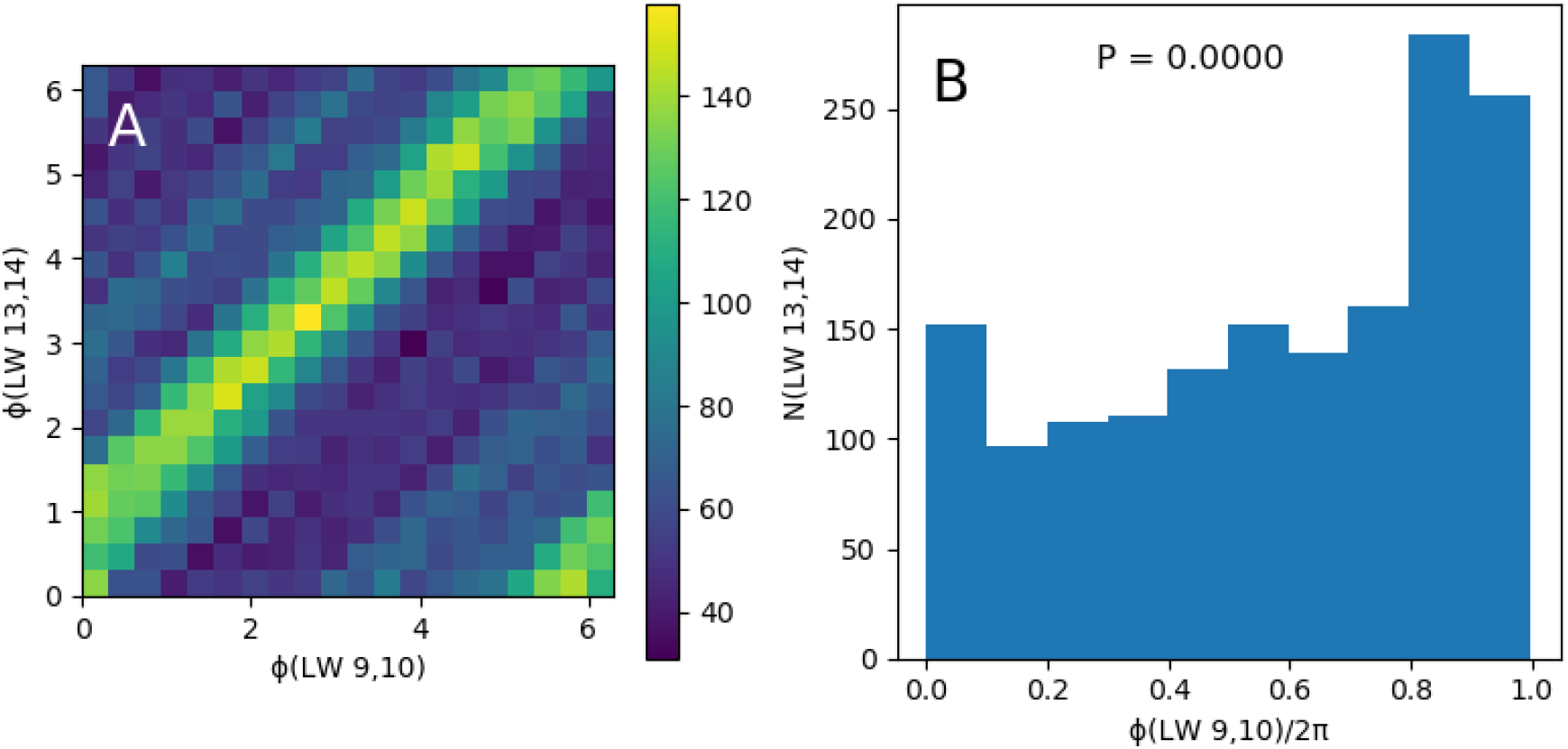
A representative recurrence map (a) and stroboscopic histogram (b). This is a high correlation between the two channels (LW 13,14 and LW 9,10) in this example.

To test whether the correlation between two phase channels is statistically significant, we tabulate the phase difference in a stroboscopic histogram. Let *φ*_1_ be the primary channel. We sample the other phase signal (*φ*_2_) whenever *φ*_1_ wraps around from 2*π* back to 0 and summarize the resulting phases in a histogram (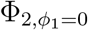 histogram) (Figure 3B). The null hypothesis (i.e., *φ*_1_ and *φ*_2_ are uncorrelated) is tested by calculating the *χ*^2^ statistics for the 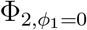 histogram, comparing it to a uniform histogram. Note that 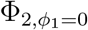 is different than 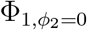.

### E. Frequency Domain Analysis

The time-domain phase-comparison methods described above are the main tools used in this paper for data analysis. However, additional information is gained by finding the peak frequency (the dominant frequency) of a group of highly correlated channels. To do so, we collect the phase signals of the channels of interest into a matrix and perform principal component analysis (PCA). Let *φ* be the principal component, i.e. the eigenvector with the largest eigenvalue. The dominant frequency is found by calculating the power spectrum of *φ* and finding its peak.

## IV. RESULTS

The recurrence maps and frequency spectra were used to deduce the number and mutual interaction of drivers during ten AF ablation procedures. Three general patterns are identified: a single driver (type I), two or more drivers with distinct frequency peaks (type II), and two or more drivers with close dominant frequencies (type III). Here, we define close dominant frequencies as a difference of less than 0.5 Hz. One patient in each of the type I and type II groups, and two patients in the type III group were on a membrane-active anti-arrhythmic medication (Vaughan-Williams class I and III) at the time of the procedure).

*Type I: Single Driver (n=3)*: Figure 4 depicts a representative case. The recurrence map for the right atrial catheter shows strong 1:1 correlations between neighboring channels. The left atrial catheter similarly captures a strong 1:1 correlation. Figure 4C is the recurrence map between the left and right atria and shows a weaker, albeit easily discernible, phase shifted 1:1 relationship. For example, Figure 4D depicts the recurrence map between two left and right atrial channels, which reaches statistical significance in the stroboscopic analysis (*P <* 0.0001, Figure 4E). Finally, the left and right dominant frequencies are the same (5.3 Hz, Figure 4F).

**FIG. 4.**
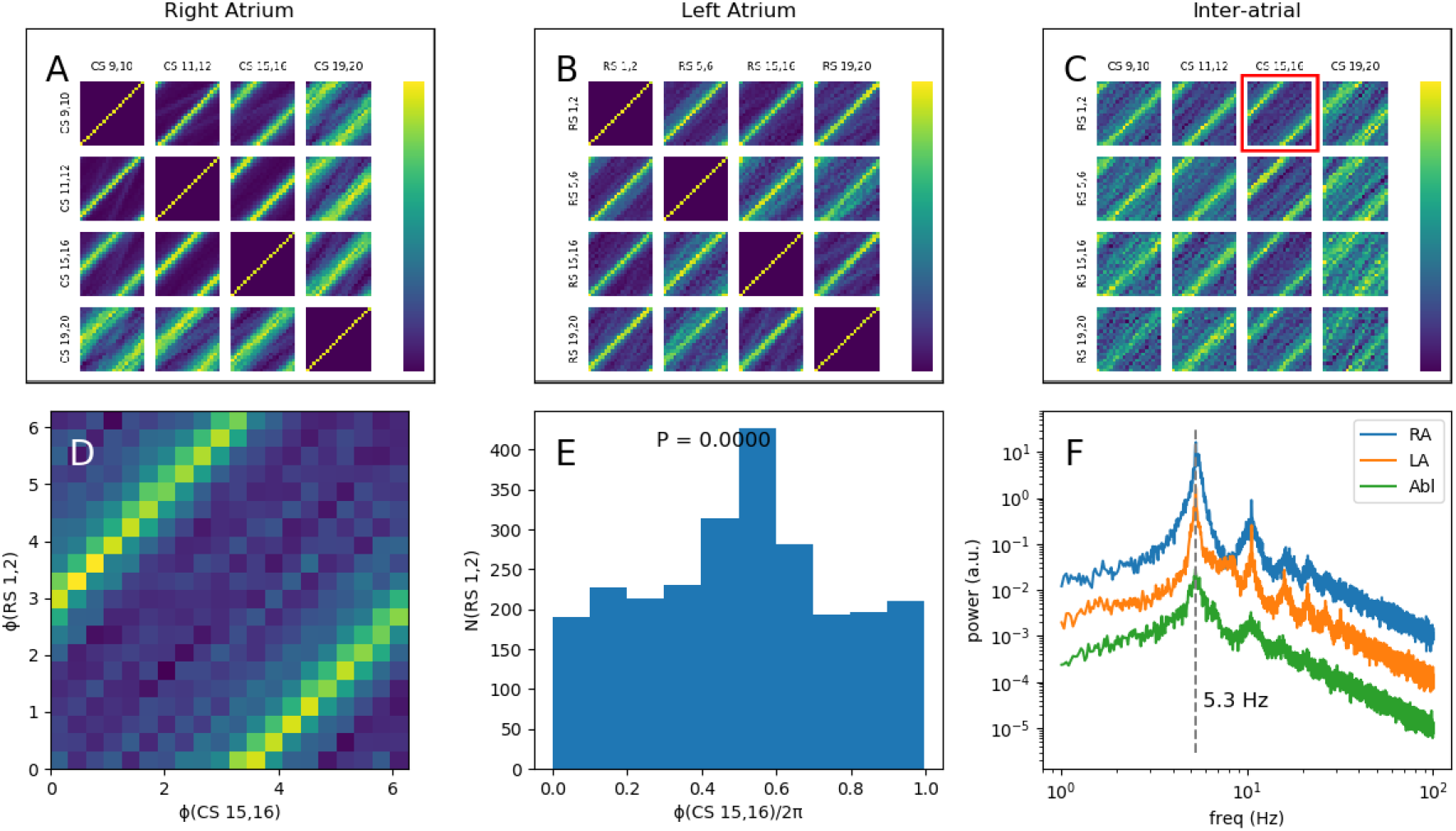
A type I example, where (a-c) depict recurrence maps for the right atrial (marked as CS, but actually from a catheter placed in the right atrium parallel to the tricuspid annulus with its tip in the coronary sinus.), left atrial (RS), and inter-atrial comparison. (d) shows the details of the recurrence map between RS 1,2 and CS 15,16 (the red square in (c)). (e) is the corresponding stroboscopic histogram to (d). (f) shows the spectra of the right atrium (blue), left atrium (orange), and the ablation catheter (green). The left atrial and ablation catheter spectra are shifted vertically to reduce clutter. Note the high intra-atrial coherence and inter-atrial correlation and similar frequencies on all channels, consistent with the presence of a single driver.

These results are indicative of the presence of a single rotor driving both atria with the same frequency. However, the variable delay between the driver and each atrium weakens the mutual correlation between them. This pattern is consistent with the “mother rotor” hypothesis.

*Type II: Two or more drivers with distinct frequencies (n=4)*: type II is the opposite of type I. In type II, each atrium is coherent, i.e., has correlated signals (Figures 5A and 5B), but the inter-atrial correlation is very poor. There is barely a suggestion of a high order regularity (2:1) in Figures 5C and 5D, which does not reach statistical significance. The dominant frequencies are also widely different (RA 5.0 Hz, LA 8.9 Hz).

**FIG. 5.**
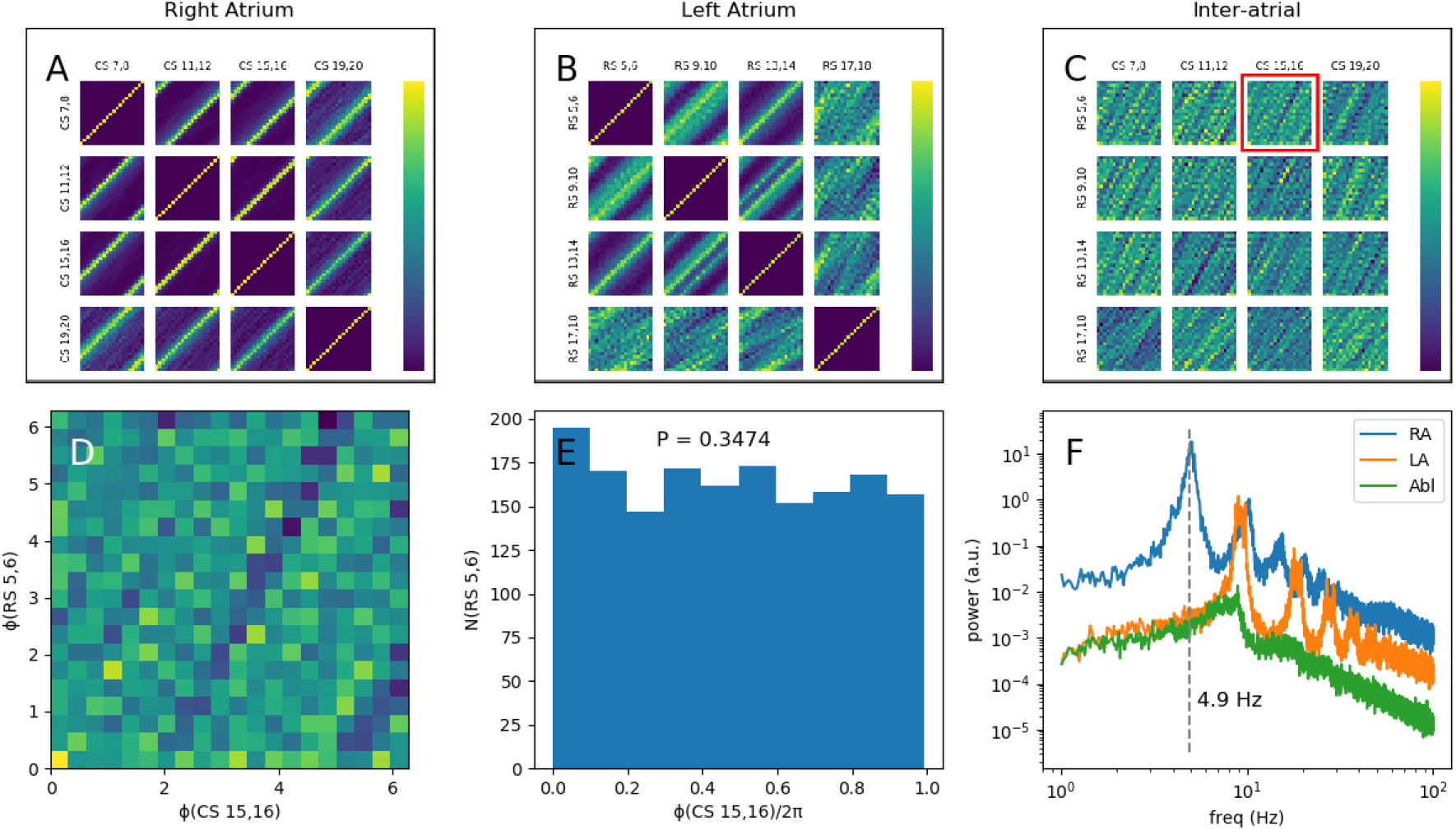
A type II example, where (a-c) depict recurrence maps for the right atrial (CS), left atrial (RS), and inter-atrial comparison. (d) shows the details of the recurrence map between RS 5,6 and LW 13,14 (the red square in (c)). (e) is the corresponding stroboscopic histogram to (d). (f) shows the spectra of the right atrium (blue), left atrium (orange), and the ablation catheter (green). The left atrial and ablation catheter spectra are shifted vertically to reduce clutter. There is high intra-atrial coherence, but very poor inter-atrial correlation. The left and right atrial peak frequencies are significantly different (the ablation catheter is located in the left atrium and has the same frequency). This pattern is consistent with the type II pattern.

These findings are consistent with the presence of at least two separate drivers, which in case of Figure 5 drive the left and right atrium separately. The large difference between the respective dominant frequencies prevents the two atria from synchronizing.

The number of drivers in type II is not limited to two. In two cases, at least three separate regions with different frequencies were observed.

*Type III: Two or more drivers with close frequencies (n=3)* type III is a composite of the other two types. There are at least two drivers with different but close frequencies (for example, 5.4 Hz and 5.8 Hz in Figure 6). There is also a statistically significant correlation between channels from the left and right atria that implies long-distance correlation. The left atrial catheter in Figure 6 is located at the antrum of the left pulmonary veins, which is 5-10 cm away from the right atrial catheter. Despite the distance, we clearly observe a 1:1 correlation between the left/right signals, which shows that the correlation length in AF can be much longer than expected.

**FIG. 6.**
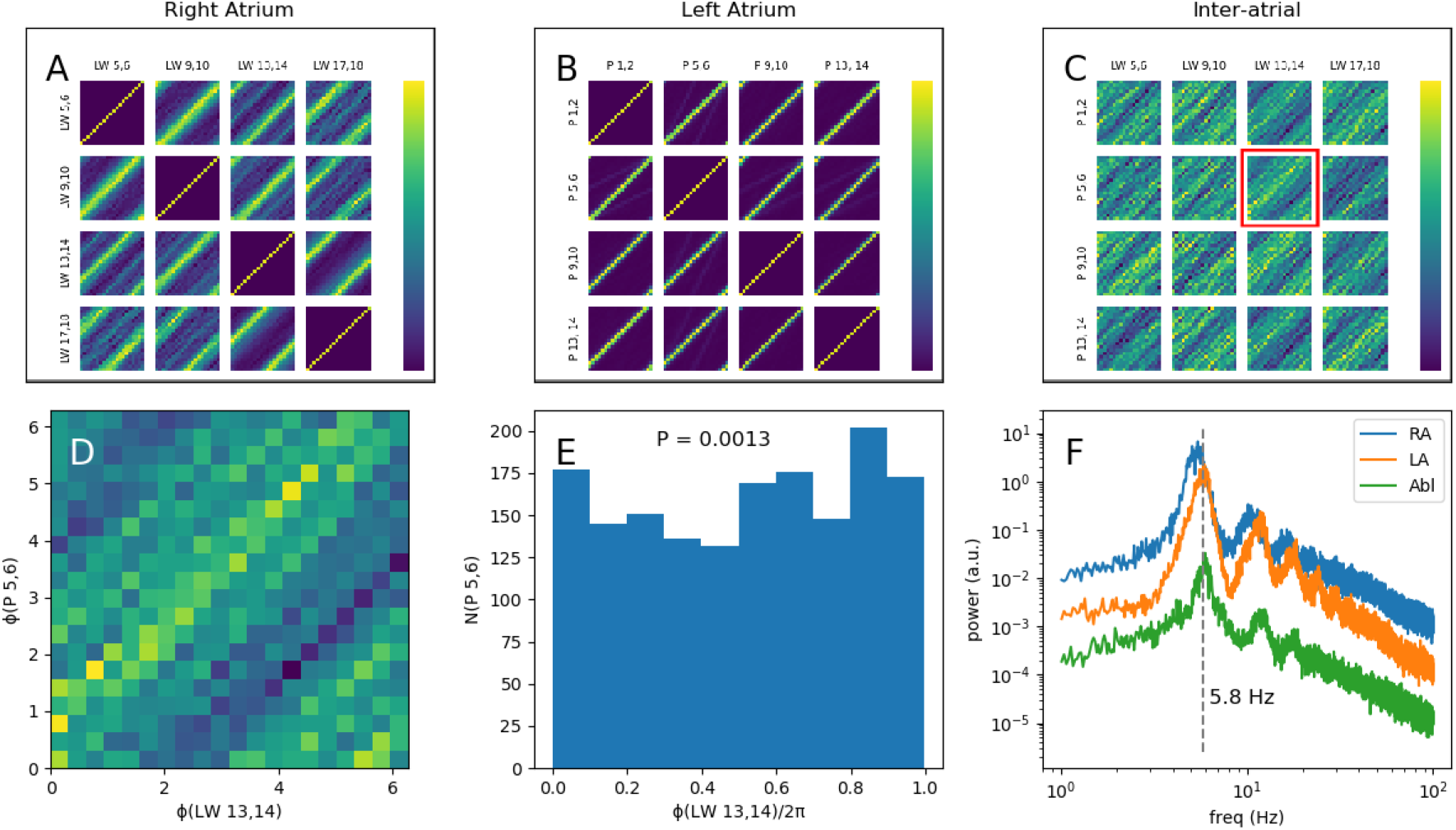
A type III example, where (a-c) depict recurrence maps for the right atrial (CS), left atrial (P), and inter-atrial comparison. (d) shows the details of the recurrence map between P 5,6 and LW 13,14 (the red square in (c)). (e) is the corresponding stroboscopic histogram to (d). (f) shows the spectra of the right atrium (blue), left atrium (orange), and the ablation catheter (green). The left atrial and ablation catheter spectra are shifted vertically to reduce clutter. There is high intra-atrial coherence, with weak but easily seen correlation between the left and right atria that reaches statistical significant in (e). The peak frequencies are slightly different (the ablation catheter is located in the left atrium and has the same frequency. This pattern is consistent with type III.

### A. Intermittency

Types I and II patterns are conceptually straightforward. In type I, there is one rotor that drivers both atria, and in type II, there are two or more separate drivers with minimum interaction. The dynamics of type III is more subtle. Now, there are at least two strongly-interacting drivers with close frequencies. We can conceptualize each rotor as a phase oscillator. Two uncoupled phase oscillators with different frequencies do not exhibit phase preference (for example, as seen in Figure 5D/E). The observed correlation between the left and right atria in type III is consistent with periods of phase locking and synchronization between the two phase oscillators. This behavior is a manifestation of *intermittent phase synchronization* or *intermittency*.^18,19^ The phases of the two rotors lock for a short time (called laminar phase periods), resulting in equal frequencies before the lock breaks and the phases start drifting apart again (turbulent phase periods), until the next time the phases lock again.

Figure 7 depicts a better example of intermittency. For this example, we are mainly interested in the dynamics of the right atrium (the left/right dynamics is of type III and similar to Figure 6). The recurrence map in Figure 7A is divided into two zones. Each zone is strongly coherent, but the correlation between the two is weak. The interesting point is the behavior of the channel between the two (CS 7,8). It is strongly correlated with both zones despite the difference between the two. The same pattern also applies to the spectra (Figure 7C). The two zones have different peak frequencies, with the CS 7,8 peak between the two. These findings are consistent with two drivers, one drives the proximal channels (CS 9,10 to CS 13,14 in the figure), and the other one controls the distal channels (CS 1,2 to CS 5,6). CS 7,8 straddles the two and becomes intermittently in phase with one and then the other. In other words, CS 7,8 resides on the dynamic border between the two zones, while the zones are driven individually by different rotors.

**FIG. 7.**
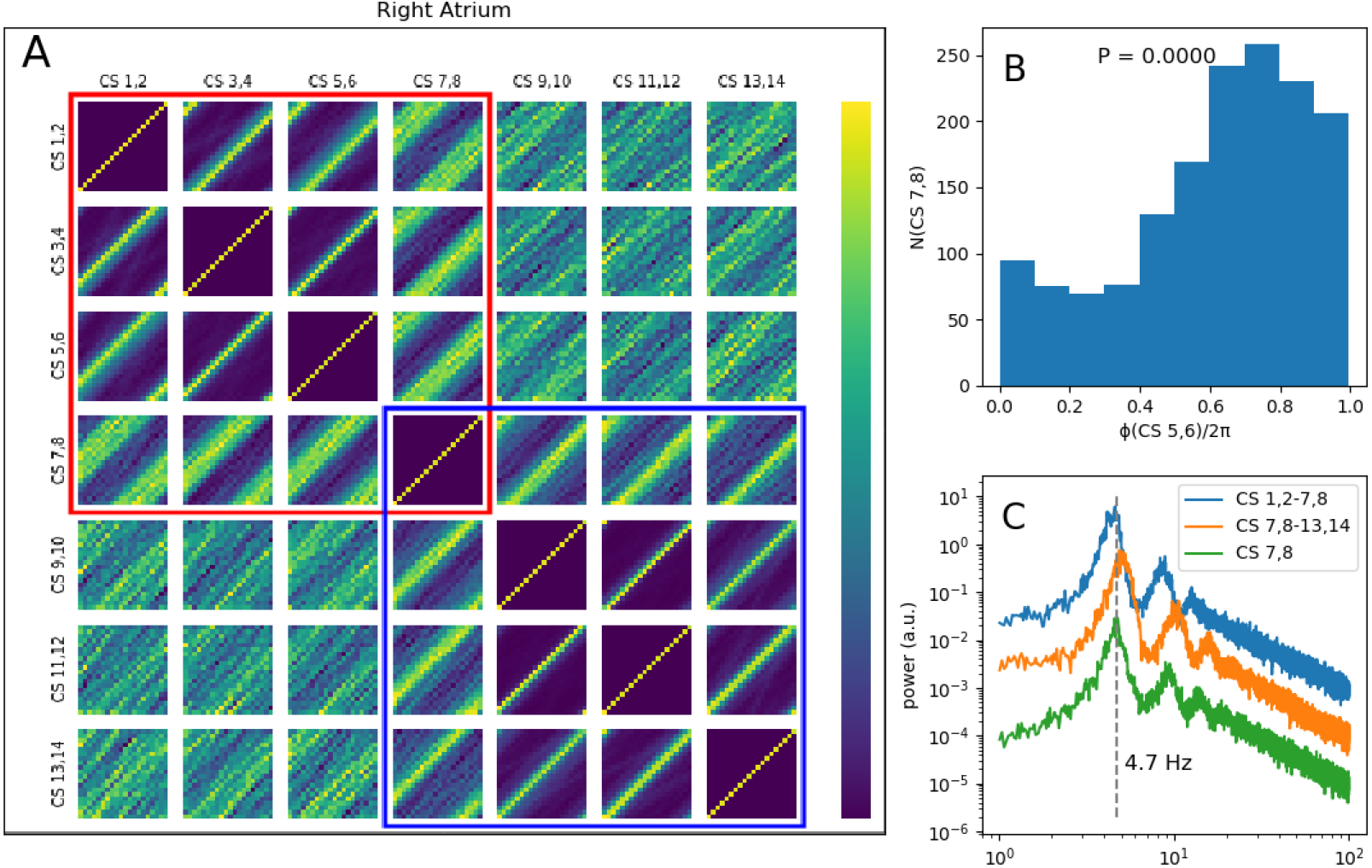
An example of intermittent phase synchronization. (a) shows the recurrence map for seven channels on the right atrial catheter. Note that there are two coherent clusters; one covers CS 1,2-CS 7,8 (the red square), and the other covers CS 7,8 - CS 13,14 (the blue square). CS 7,8 resides in both clusters (see the text for details). (b) depicts the histogram between CS 7,8 and CS 5,6. The two clusters have different frequencies with the shared channel (CS 7,8) in between (c).

## V. DISCUSSION

The analysis of intracardiac signals recorded during AF ablation procedures provides mechanistic insight into the dynamics of AF. The analytic methodology described in the Methods section – the recurrence maps and dominant frequency analysis – can detect and quantify subtle correlations between different atrial sites.

AF is a chronic, progressive, and heterogeneous disease, with multiple mechanisms and variable dynamics among different patients or longitudinally in time in the same person. The complexity of AF and the lack of proper animal models render a detailed mechanistic analysis very difficult.^15^ An ideal dataset should have complete spatial coverage for an extended period. Non-invasive methods, including body-surface multi-electrode techniques based on solutions to the inverse problem,^20^ have a low effective spatial resolution. Invasive methods, based on recordings during electrophysiology procedures or cardiac surgeries, have practical and ethical restrictions and are limited to short segments from a few points. In this paper, a set of analytic tools was developed to circumvent these limitations. These tools are based on the framework of the synchronization theory and allow us to describe AF and obtain meaningful information out of the incomplete and noisy recordings.^18,19,21^

Atrial activity in AF is classified into three main types (Figure 8). Type I is consistent with the mother rotor hypothesis with one dominant rotor driving both atria. The majority of the cases have more than one driver and are of types II and III. Previous studies have shown that AF is organized into interacting areas,^9,22^ which may shrink, enlarge, or even merge during ablation and are similar to the driver regions in this paper.

**FIG. 8.**
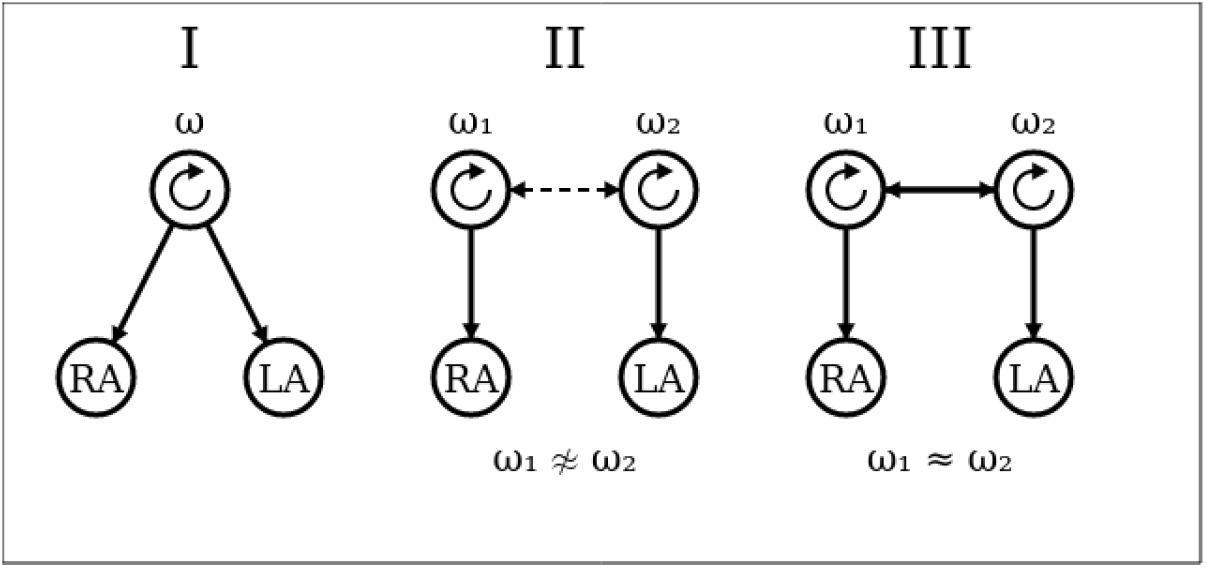
The schematic diagram of the three atrial organizational types.

The key difference between types II and III is the mutual interaction between drivers. Each rotor (spiral wave) is akin to a phase oscillator. Additionally, spiral waves can have an excitable gap.^23^ Therefore, for a specific phase difference between two interacting spiral waves, the activation wavefront from one may enter the excitable gap of the second one and reset its phase. Naively, we would expect the faster rotor to eventually entrain the slower one. However, in contrast to the situation in atrial flutter, this does not happen in AF. Rotors in AF are operating near the conduction capacity of the atrial tissue, such that complete 1:1 entrainment is not possible. Instead, the slower rotor only intermittently feels the effect of the faster one. The lack of complete entrainment protects the slower one and allows it to also intermittently affect the faster one. In the language of synchronization, we are dealing with two or more weakly coupled oscillators. If the intrinsic frequencies are close (type III), the coupling is constructive and facilitates 1:1 entertainment. However, when the intrinsic frequencies are far apart (type II), the effect of coupling can be similar to noise and may contribute to the irregular nature of AF.

This last point has practical implications. Membrane-active anti-arrhythmic medications and ablation reduce coupling and may potentially suppress noise, which can lead to partial regularization of the rhythm. Moreover, the type of AF may determine its response to ablation. For example, it is plausible that type III AF is more likely than type II to regularize and convert to flutter due to ablation.

Another finding was the existence of long-distance correlation in AF, where detectable 1:1 correlation between points physically far apart (e.g., lateral right atrium to the left pulmonary veins) was observed. The correlation was weak and not apparent visually or using standard analytic methods. Such long-distance correlations were not limited to type I but were also seen in type III with multiple drivers. Hence, AF is more organized than expected based on the irregular and seemingly chaotic atrial signals. It is plausible that the magnitude of the underlying organization modulates the success of ablation.

## Notes

### Competing Interest Statement

The authors have declared no competing interest.

